# High Tau Expression Correlates with Reduced Invasion and Prolonged Survival in Ewing Sarcoma

**DOI:** 10.1101/2025.02.10.635652

**Authors:** Florencia Cidre-Aranaz, Claudia Magrin, Malenka Zimmermann, Jing Li, Anna Lisa Baffa, Matteo Ciccaldo, Wolfgang Hartmann, Uta Dirksen, Martina Sola, Paolo Paganetti, Thomas G P Grünewald, Stéphanie Papin

**Affiliations:** Hopp-Children’s Cancer Center (KiTZ), Heidelberg, Germany; Division of Translational Pediatric Sarcoma Research, German Cancer Research Center (DKFZ), German Cancer Consortium (DKTK), Heidelberg, Germany; National Center for Tumor Diseases (NCT), NCT Heidelberg, a partnership between DKFZ and Heidelberg University Hospital, Germany; Laboratory for Aging Disorders, Laboratories for Translational Research, Ente Ospedaliero Cantonale, Bellinzona, Switzerland; Faculty of Biomedical Sciences, Università della Svizzera Italiana, Lugano, Switzerland; Gerhard-Domagk-Institute of Pathology, Münster University Hospital, Münster, Germany; Pediatrics III, University Hospital Essen, West German Cancer Center, German Cancer Consortium (DKTK) site Essen, National Center for Tumor Diseases (NCT) West, Essen, Germany; Faculty of Medicine, Heidelberg University, Heidelberg, Germany; Institute of Pathology, Heidelberg University Hospital, Heidelberg, Germany

**Author notes:** co-corresponding authors: Prof. Dr. Paolo Paganetti, Laboratories for Translational Research, via Chiesa 5, CH-6500 Bellinzona, Switzerland, Prof. Dr. Dr. Thomas G P Grünewald, Division of Translational Pediatric Sarcoma Research, German Cancer Research Center (DKFZ), Im Neuenheimer Feld 280, 69120 Heidelberg, Germany. co-first authors (Florencia Cidre-Aranaz, Claudia Magrin). co-senior authors (Paolo Paganetti, Thomas G P Grünewald, Stéphanie Papin).

## Abstract

The microtubule-associated protein Tau (encoded by the *MAPT* gene) is linked to a family of neurodegenerative disorders defined as tauopathies, which are characterized by its brain accumulation in neurofibrillary tangles and neuropil threads. Newly described Tau functions comprise DNA protection, chromatin remodeling, p53 regulation and cell fate modulation, suggesting a role of Tau in oncogenesis. Bioinformatic-supported characterization of Tau in cancer reveals robust expression in bone cancer cells, in particular Ewing sarcoma (EwS) cell lines. EwS is an aggressive cancer caused by a fusion of members of the *FET* and *ETS* gene families, primarily *EWSR1::FLI1*. Here we found that *MAPT* is a EWSR1::ETS target gene and that higher Tau expression in EwS cells inhibited their migratory and invasive behavior, consistent with a more immobile and proliferative phenotype observed in EwS. Indeed, we report that high Tau expression is associated with improved overall survival of EwS patients. We also show that the sessile but proliferative phenotype of EWSR1::ETS-high cells may result from a modulatory role of Tau on focal adhesion to extracellular matrix proteins. Our data highlight the utility of determining Tau expression as a prognostic factor in EwS as well as the opportunity to target Tau expression as an innovative EwS therapy.

## Introduction

Tauopathies comprise a group of neurodegenerative disorders including Alzheimer’s disease (AD) and are characterized by progressive Tau deposition in neurofibrillary tangles^1^. Autosomal dominant mutations in *MAPT* encoding for microtubule-associated Tau proteins are responsible for frontotemporal lobar degenerations (FTLD-Tau)^2–4^. A prevalent hypothesis for the pathogenesis of tauopathies is that aberrant Tau modification, such as increased phosphorylation and a rigid conformation, contributes to Tau dissociation from microtubules and multimeric β-sheet assembly, which correlates with proteotoxicity, neuronal dysfunction and cell death^5, 6^.

Interestingly, non-canonical functions have been reported for Tau. For instance, Tau translocates to the cell nucleus upon heat or oxidative stress and protects DNA^7–11^. Neurons knocked-out for *MAPT* display enhanced DNA damage^12^, and induced DNA damage correlates with nuclear translocation and dephosphorylation of Tau^13^. Chromosomal abnormalities in AD fibroblasts^14^ and frequent DNA damage in AD brains^15, 16^ both reinforce the emerging Tau function in DNA stability. Tau depletion also deregulates DNA damage-induced apoptosis and cell senescence by a mechanism involving MDM2-dependent p53 degradation^17, 18^. Additional functions of Tau in epigenetic modulation were reported^19, 20^. Upon histone binding, Tau stabilizes chromatin compaction^19, 21–24^ and affects global gene expression during the neurodegenerative process^21, 22^.

All these non-canonical functions hint to an implication of Tau in cancer. In fact, several studies reported a correlation between *MAPT* mRNA expression and survival in various cancer types^25–27^. Whereas a positive correlation is found in neuroblastoma, breast cancer, low grade glioma, kidney clear cell carcinoma, lung adenocarcinoma and pheochromocytoma-paraganglioma, a negative correlation is reported in colon cancer and head and neck cancers, liver hepatocellular carcinoma and uterine corpus endometrial carcinoma^25, 27–29^. A recent bioinformatic study suggests that the implication of Tau in cancer is likely mediated by modulation of three main pathways: epithelial-to-mesenchymal transition (EMT), inflammation, and regulation of the cell cycle^27^. In addition, Tau mutations are associated to an increased cancer risk^30^. Microtubule-related and non-canonical functions of Tau may underlie these observations^17, 20, 26^.

Here we report that Tau is highly expressed in Ewing sarcoma (EwS) – an aggressive bone or soft tissue cancer mainly affecting children, adolescents, and young adults. EwS is caused by a chromosomal translocation generating a fusion protein with aberrant transcriptional activity composed of members of the FET and ETS gene families, primarily EWSR1::FLI1^31^. We demonstrated that *MAPT* mRNA expression in EwS tumors positively correlates with survival, possibly mediated by a modulatory role of Tau on EwS cell adhesion, migration, and invasion.

## Results

### *MAPT* is a target gene of the EWSR1::FLI1 fusion protein

To first characterize the expression of Tau in different cancer-derived cell lines, we mined the DepMap data collection (https://depmap.org/portal/). As expected, neuroblastoma lines harbored the highest *MAPT* gene expression. Strikingly, bone-derived cell lines showed the second strongest *MAPT* expression, while portraying a strong inter-cell line variability (**Fig. 1A**). To better understand this aspect, we explored the *MAPT* expression in each different bone sarcoma, and observed that EwS cell lines presented a significantly higher *MAPT* expression than osteosarcoma and chondrosarcoma cell lines (**Fig. 1B**). These results were further validated using gene expression microarray data derived from a distinct panel of 18 EwS cell lines expressing EWSR1::FLI1 (type 1 or type 2) or EWSR1::ERG^32^. Here again, most cell lines, beside A-673 and CHLA-25, displayed high *MAPT* transcripts (**Table 1**).

**Figure 1.**
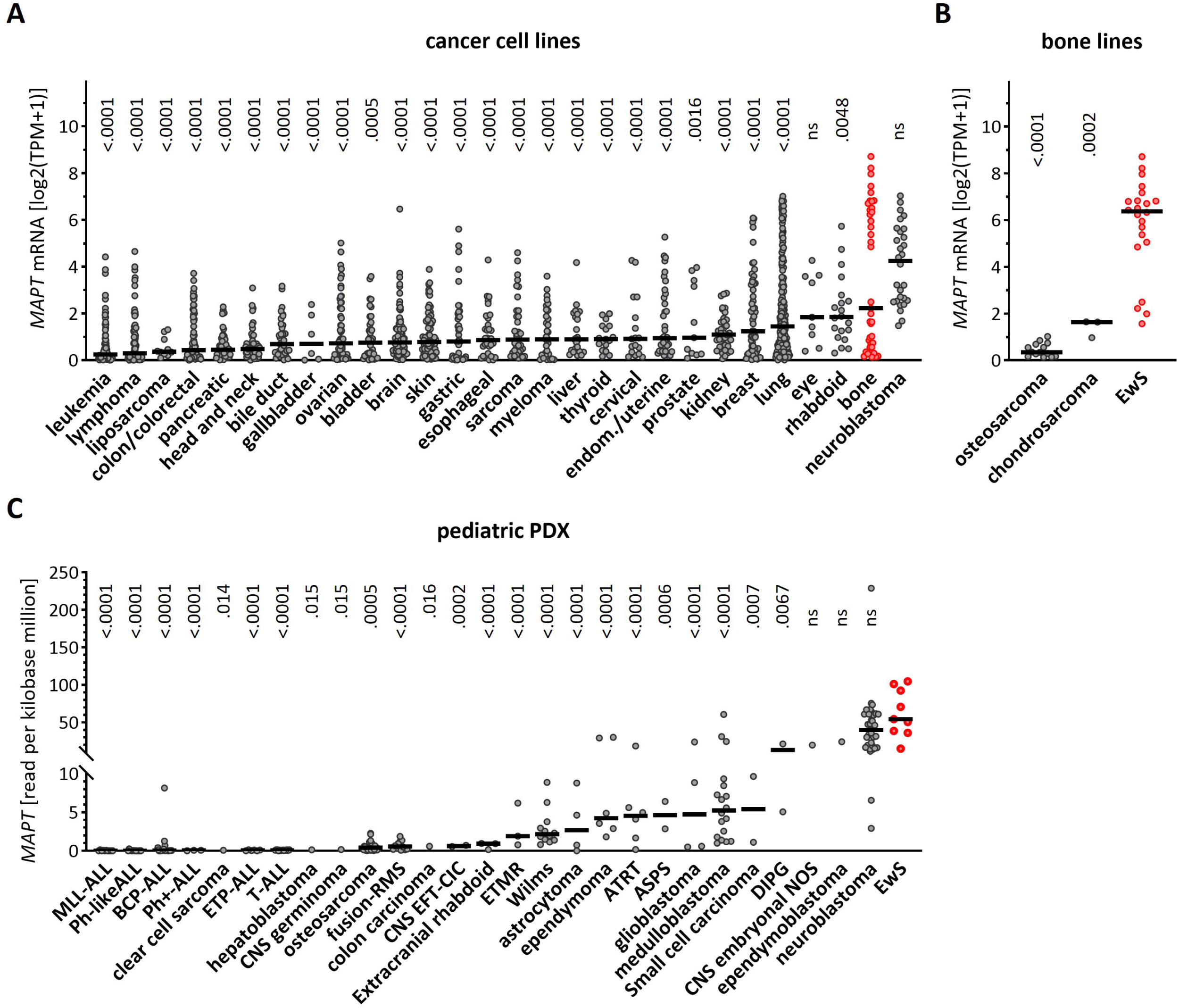
MAPT mRNA is highly expressed in EwS. **A.** MAPT transcript in DepMap cell lines originating from the indicated cancer types. TPM = transcript per million. Horizontal bar = median. One-way ANOVA (p<0.0001) and Dunnett’s multiple comparison to EwS (in red). **B.** MAPT mRNA in bone-derived EwS, osteosarcoma and chondrosarcoma. Horizontal bar = median. One-way ANOVA (p<0.0001) and Tukey’s multiple comparison to EwS (in red). **C.** MAPT mRNA in patient-derived mouse xenograft derived from the indicated cancer types. Horizontal bar = median. One-way ANOVA (p<0.0001) and Dunnett’s multiple comparison to EwS (in red).

We then analyzed *MAPT* expression in patient-derived xenografts (PDX) from the pediatric preclinical testing consortium^33^. In agreement with our previous observations, EwS PDXs displayed the highest *MAPT* mRNA levels as compared to all other pediatric tumors (**Fig. 1C**). Thus, EwS cells were characterized by high *MAPT* gene expression.

To better characterize the potential interplay between *MAPT* and EWSR1::FLI1, we first analyzed single cell transcriptomic data derived from three EwS PDX in comparison with two primary cultures of mesenchymal stem cells (MSC)^34^, a proposed EwS cell-of-origin^35^. Interestingly, the MSC primary cultures showed negligible levels of *MAPT* mRNA, whereas the three EwS PDXs showed significantly higher *MAPT* expression (**Fig. 2A**), thus suggesting a potential regulatory role of the EwS fusion protein on *MAPT* expression. To further explore this interplay, we analyzed data derived from 18 EwS cell lines with a doxycycline-inducible EWSR1::ETS-knock-down (KD)^32^. In the great majority of the lines, EWSR1::FLI1 or EWSR1::ERG-KD reduced *MAPT* transcripts (**Fig. 2B**) and Tau proteins (**Fig. 2C**). ChIP-Seq data from these EwS cell lines (http://r2platform.com/escla/) revealed a prominent EWS::ETS peak close to the *MAPT* gene corresponding to a core EWS::ETS site^32^ in all cell lines analyzed (**Fig. 2D**). In agreement with the induction of H3K27 acetylation upon EwS fusion proteins binding to DNA^32^, a concomitant H3K27Ac peak was also detected. Overall, these results indicated that EWSR1::FLI1 and EWSR1::ERG induced *MAPT* expression.

**Figure 2.**
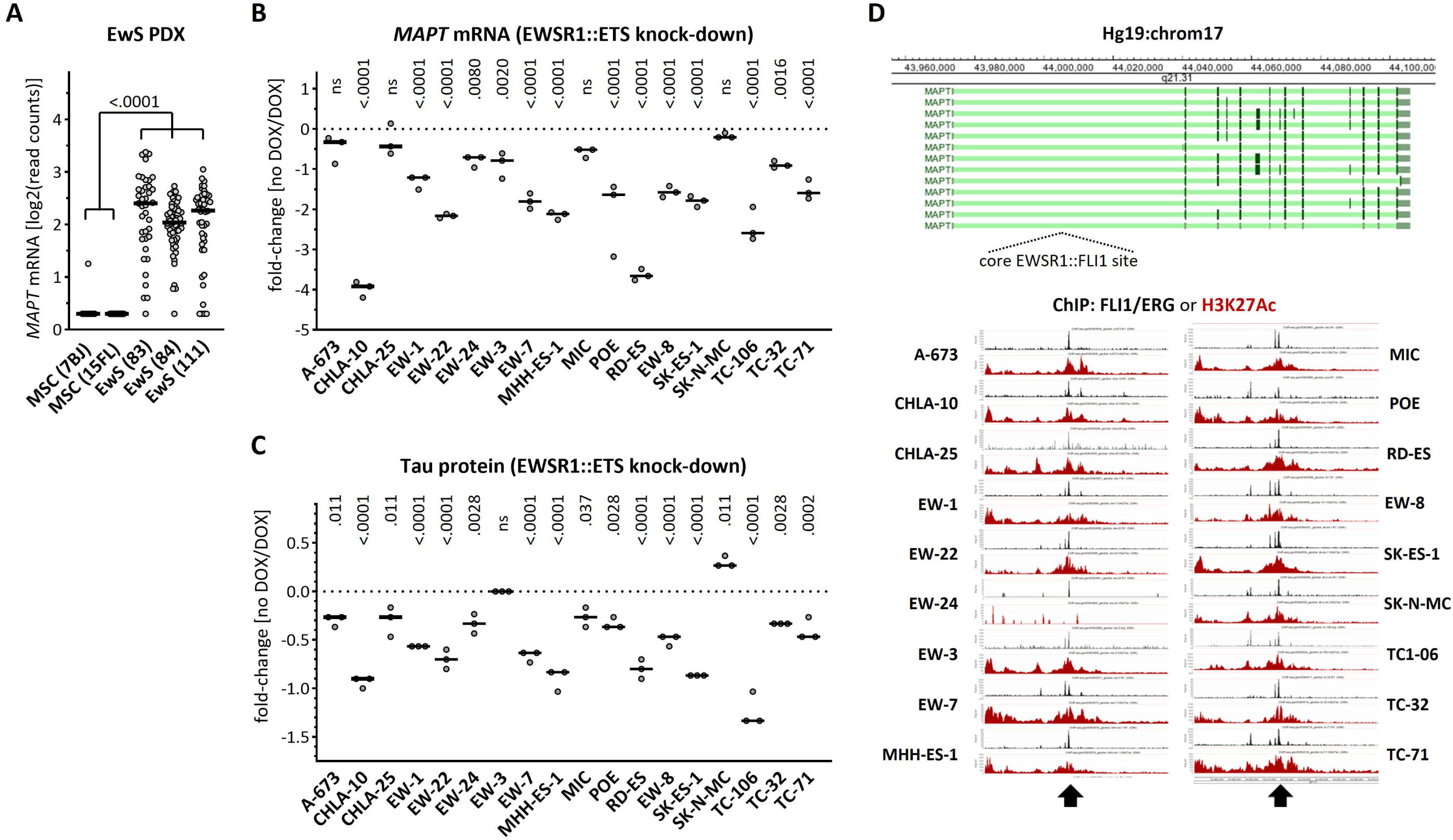
MAPT is a target gene of the EWSR1::FLI1 fusion protein. **A.** Single cell MAPT mRNA in EwS patient-derived mouse xenografts or in mesenchymal stem cells (MSC). Horizontal bar = median. One-way ANOVA (p<0.0001) and Tukey’s multiple comparison. **B.** Relative expression of MAPT transcripts in the indicated EwS cell lines upon doxycycline-induced downregulation of the EWSR1::FLI1 fusion protein normalized for non-induced cells. 2way ANOVA (p<0.0001 for cell line and knock-down) and Šídák’s multiple comparison between non-induced and induced for each cell line. **C.** Relative expression of Tau protein in the indicated EwS cell lines upon induced downregulation of the EWSR1::FLI1 fusion protein normalized for non-induced cells. 2way ANOVA (p<0.0001 for cell line and knock-down) and Šídák’s multiple comparison between non-induced and induced for each cell line. **D.** Identification of chromatin immunoprecipitation events along the MAPT gene corresponding to EWSR1::FLI1 and H3K27ac binding in the indicated EwS cell lines.

### Tau-KD affects EwS cell adhesion, migration, and invasion

To explore the role of the EWSR1::FLI1-dependent increase in Tau expression for cancer-associated cellular phenotypes, we engineered the EwS cell line TC-32 with inducible inhibition of Tau expression (**Fig. 3A)** and the EwS cell line TC-71 with constitutive inhibition of Tau expression (**Fig. 3B**). For the TC-32 line in particular, Tau downregulation was relatively modest after induction of the Tau shRNA for at least two weeks, perhaps because of the long life of Tau protein^36^. Nonetheless, under these conditions of partial Tau-KD, we observed decreased cell proliferation for both lines (**Fig. 3C-D**), an observation conceivably linked to the microtubule-binding function of Tau during mitosis^37, 38^.

**Figure 3.**
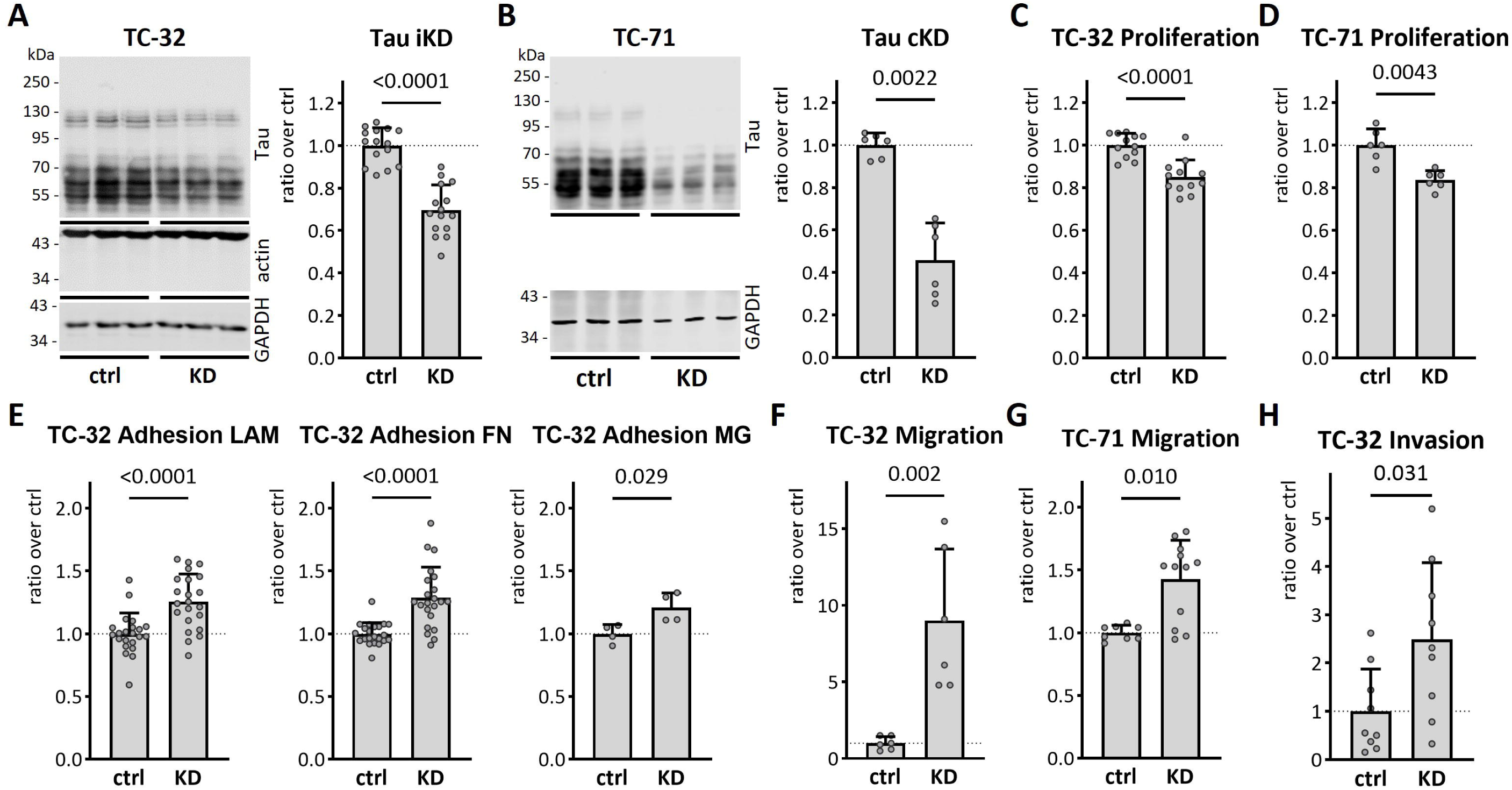
Tau protein knock-down affects adhesion, migration, and invasion. **A.** Lysates from biological triplicates from inducible TC-32 Tau-KD cells not (ctrl) or treated (KD) with doxycycline for 2.5 weeks were analyzed by western blot with the mouse Tau13 antibody, the mouse actin antibody, and the rabbit GAPDH antibody, followed by visualization with anti-mouse IgG IRDye 680RD and anti-rabbit IgG IRDye 800CW. Intensity of the Tau and GAPDH signals is reported as ratio Tau/GAPDH (n=15). **B.** Lysates from biological triplicates were prepared from constitutive TC-71 Tau-KD and analyzed by western blot with Tau13 and GAPDH antibodies visualized with anti-mouse IgG IRDye 680RD and anti-rabbit IgG IRDye 800CW. Signal intensity is reported as ratio Tau/GAPDH (n=6). **C.** TC-32 Tau-KD cells in MW96 were imaged for confluency at the Incucyte every 2 h and cell proliferation calculated over 72 h starting from time 0 (n=12). **D.** Same as in C. for TC-71 Tau-KD cells (n=6). **E.** Adherent TC-32 Tau-KD cells 30 min after plating on laminin (LAM), fibronectrin (FN) or Matrigel (MG) were stained with crystal violet (n=22 for LAM and FN, n=4 for MG). **F.** TC-32 Tau-KD cells were seeded in transwell inserts without any coating. Cells were imaged with crystal violet on the other side of the filters 2 days later and quantified with ImageJ (n=6). KD data were normalized over ctrl, reported as mean ± SD and analyzed with the Mann-Whitney test. **G.** As in F. for TC-32 Tau-KD cells seeded in transwell inserts pre-coated with Matrigel. **H.** A wound was automatically generated on a TC-71 Tau-KD cell monolayer in MW96 and closure of the wound was imaged at the Incucyte every 2 h to determine their migration capabilities (n=12). KD data were normalized over ctrl, reported as mean ± SD and analyzed with the Mann-Whitney test.

It is well-accepted that metastasis is a critical factor contributing to cancer malignancy in addition to tumor growth at the primary site^39^. Metastasis may be modelled *in vitro* by testing cell adhesion, migration and invasion. Intriguingly, TC-32 cell adhesion was found increased by Tau-KD on culture plates coated with the extracellular matrix proteins laminin, fibronectin and matrigel (**Fig. 3E**). Tau-KD also increased migration of TC-32 and TC-71 EwS cells (**Fig. 3F-G**) and enhanced TC-32 cell invasion in a transwell assay (**Fig. 3H**). Concordantly, doxycycline-treatment of TC-32 control cells did not affect these phenotypes (**Suppl. Fig. 1**). This excluded a nonspecific effect of doxycycline but confirmed the participation of Tau in these cellular mechanisms. Overall, the data obtained showed that high Tau expression in EwS results in an inhibition of the migratory/invasive phenotype in EwS cell lines.

### Changes in EMT markers and morphology in Tau-KD cells

Recent studies reported a possible link between Tau expression and the EMT pathway in cancer^27^. To question the implication of this pathway in the increased adhesion, migration and invasion observed in Tau-KD cells, we determined the phosphorylation status of the focal adhesion kinase (FAK) as a key regulator of these processes^40^, and vimentin as a representative EMT marker^41^. Tau-KD increased both Tyr_397_ phosphorylation of FAK (P-FAK) and the expression of vimentin (**Fig. 4A**). No changes in P-FAK and vimentin were found in doxycycline-treated control TC-32 (**Suppl. Fig. 2**). Immune staining analysis confirmed increased P-FAK in Tau-KD cells (**Fig. 4B**). Whilst the number of focal adhesion foci visualized by phalloidin-staining was unaffected by the expression of Tau, Tau-KD cells displayed qualitatively larger focal adhesion points. In fact, Tau-KD led to more flat, elongated EwS cells displaying a prominent actin network when compared to the characteristic small and round morphology with relatively few actin filaments observed in normal EwS cells (**Fig. 4B**).

**Figure 4.**
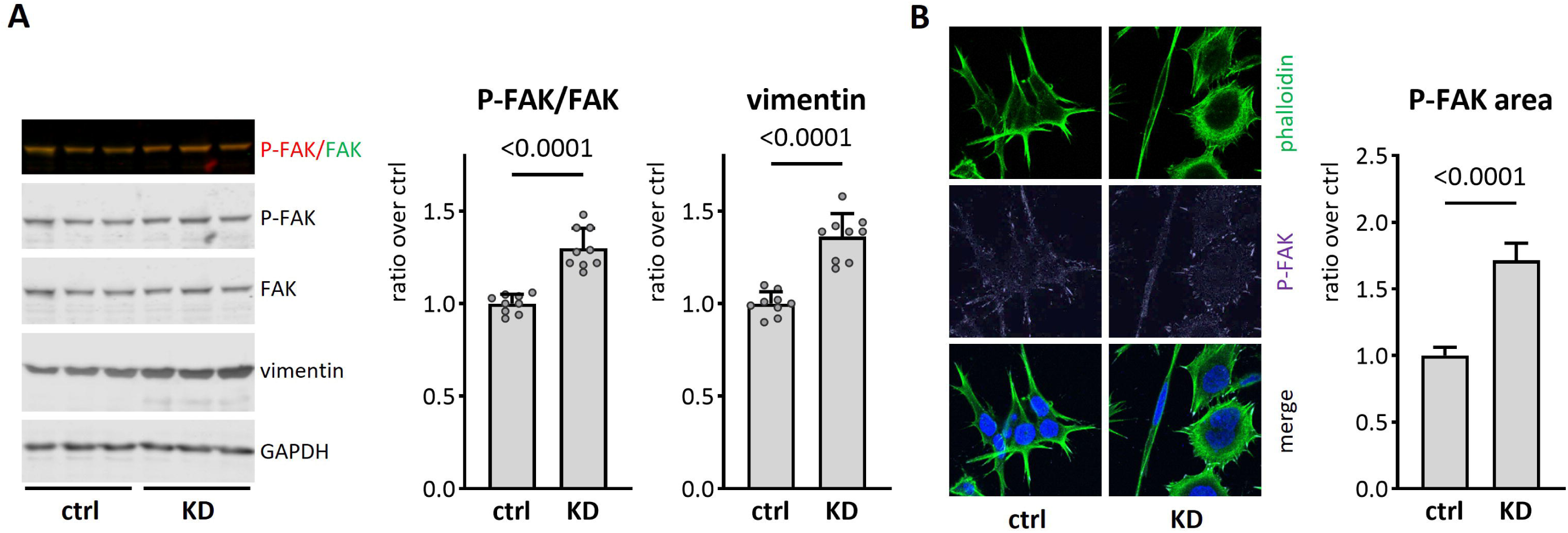
Tau-KD leads to change in EMT-like markers and cell morphology. **A.** Lysates from biological triplicates were prepared from TC-32 Tau-KD not (ctrl) or treated (KD) with doxycycline for 2.5 weeks. Samples were analyzed by western blot with P-FAK, FAK, vimentin and GAPDH antibodies. Primary antibodies were detected with anti-rabbit IgG IRDye 800CW or anti-mouse IgG IRDye 680RD. The intensity of the signal is reported as ratio P-FAK/FAK and vimentin/GAPDH (n=15). **B.** TC-32 Tau-KD cells were fixed and stained with phalloidin-AF488 and P-FAK antibody followed by secondary anti-rabbit-AF647 and counterstained with DAPI. Images were acquired on a confocal microscope and mean ± sem P-FAK focal adhesion area was quantified with ImageJ (n=738-746 adhesion points). KD data were normalized over ctrl, reported as mean ± SD and analyzed with the Mann-Whitney test.

### *MAPT* gene expression positively correlates with overall survival of EwS patients

Since our *in vitro* data suggested that high Tau expression could attenuate the aggressiveness of EwS cells by decreasing their migratory/invasive potential, we next analyzed a patient cohort composed of 166 EwS patients^32, 42^. In agreement with our *in vitro* results, this analysis revealed that high *MAPT* expression was significantly associated with better survival (*p*=0.0002) (**Fig. 5A**). To validate these results in a second cohort using a complementary method, we immunohistochemically evaluated Tau protein expression in an EwS tissue microarray containing 50 patient samples. Survival analysis on these samples confirmed the association high Tau expression with better survival in EwS (**Fig. 5B**). Collectively, these results on the association of high *MAPT* expression with better prognosis suggest a protective role of Tau in EwS.

**Figure 5.**
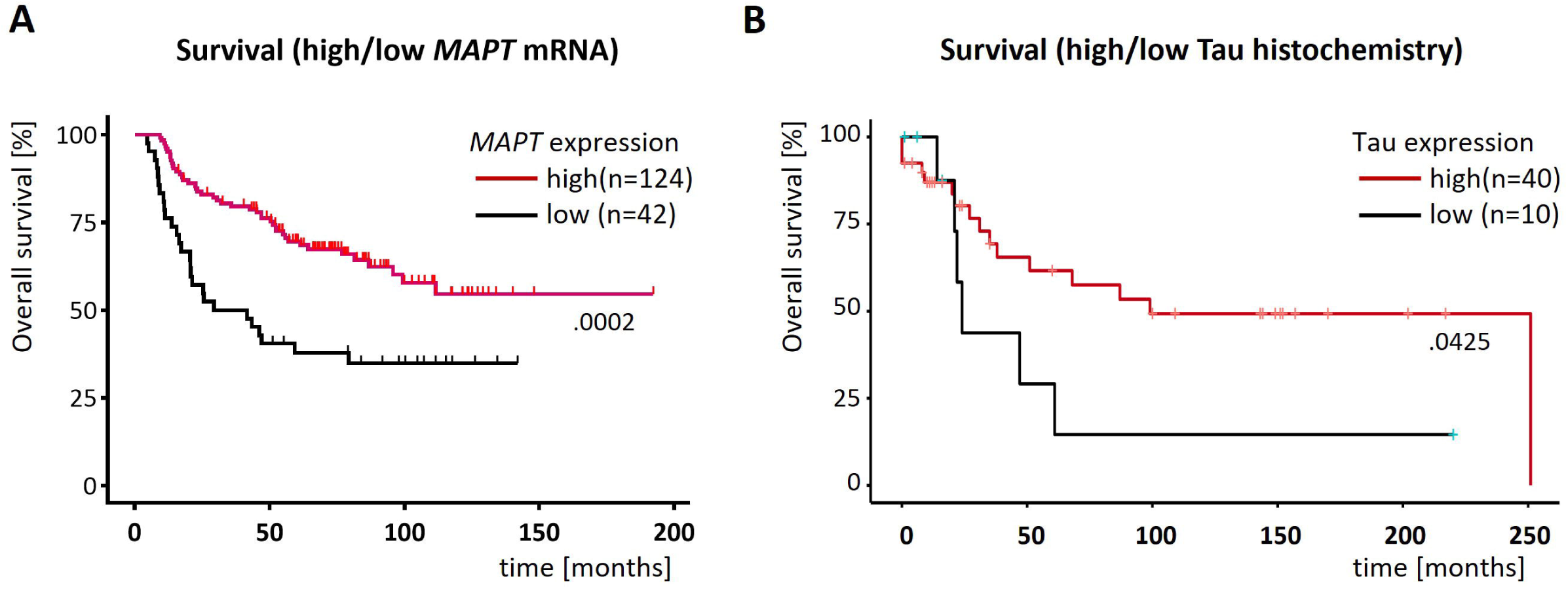
MAPT mRNA and Tau protein expression in EwS is linked to survival. **A.** Kaplan-Meier survival curve of an EwS cohort separated on the basis of expression of the MAPT mRNA in the two subgroups high (above mean value) or low (below mean value). Two-sided Log-rank (Mantel-Cox) test. **B.** as in **A.** but based on Tau protein expression determined by immunohistochemistry. One-sided Log-rank (Mantel-Cox) test.

## Discussion

We report that in EwS, *MAPT* is targeted and activated by the aberrant transcription factors EWSR1::ETS. Our data support an active involvement of the *MAPT* transcript Tau protein in the pathogenesis of EwS. *In vitro*, high Tau protein in EwS cells was associated with reduced adhesion, migration, invasion, and decreased EMT markers. Concordantly, in clinical samples EwS patient survival positively correlated with *MAPT* expression.

A positive correlation between *MAPT* mRNA levels and survival was reported also in other cancers such as low-grade glioma, breast cancer, kidney clear cell carcinoma, lung adenocarcinoma, and pheochromocytoma/paraganglioma^27, 43^. In contrast, Tau expression negatively correlates with survival in colorectal adenocarcinoma and skin melanoma^27, 43^. Among the Tau-associated genes linked to the correlation between Tau expression and patient’s survival^27^, several encode for structural proteins and extracellular matrix components, which are involved in cell adhesion, migration, and the EMT^44, 45^. A direct role of Tau in its positive correlation with survival has been explored in different glioma models^46^. In human glioblastoma LN229 cells, Tau overexpression impaired cell migration and invasion without altering cellular proliferation^47^. In contrast, Tau-KD in glioblastoma U87 cells increased Rho/ROCK signaling and inhibited cell migration^48, 49^. The full understanding of the Tau-dependent mechanisms mediating migration and invasion requires further investigation. A strong clue comes from the regulatory function of the microtubule-associated protein Tau on the organization of microtubules (MT) and the cytoskeleton^50^. Cell motility and mitosis are MT-orchestrated processes. It is therefore likely that Tau-dependent remodeling of the cytoskeleton impacts on disease in EwS, which is a highly metastatic cancer associated with exacerbated cell migration, as reported in glioma cells^46, 48, 49^.

Compellingly, the increase in migration and invasion in EwS cells with Tau-KD resembles the phenotype of EwS cells with low EWSR1::FLI1 expression. These correlate with a more aggressive metastasis spread when compared to cells with high EWSR1::FLI1 expression, that in turn are characterized by highly active proliferation but reduced invasiveness^51^. In fact, decreased EWSR1::FLI1 expression leads to major changes in effectors of actin cytoskeleton and adhesion process dynamics with a shift from cell-to-cell to cell-matrix adhesion^51^. Reciprocally, analysis of focal adhesion architecture and actin cytoskeletal organization revealed that cells expressing high level of EWSR1::FLI1 present with a loss of actin stress fibers and reduced size and number of focal adhesions, structural changes that likely account for the functional deficit in cell adhesion, an important step in the final colonization of metastatic cells^52^. Corresponding changes in cellular morphology have also been observed following expression of EWSR1::FLI1 in human mesenchymal stem cells, which causes the conversion of a well-spread, fibroblast-like shape to a rounded cell morphology^52^. In summary, these changes in the case of a low EWSR1:FLI1 expression are associated with a dramatic increase of *in vivo* cell migration and invasion potential^51^ and it is believed that this cell-to-cell variation of EWSR1::FLI1 expression levels may constitute a critical component of the early metastatic process occurring in EwS^52^. It is important to note that classical models of metastasis are derived from study of epithelial-derived tumor cells, which undergo an EMT during acquisition of metastatic potential. Non-epithelial tumors, such as sarcomas, may follow a different metastasis process. Sarcomas are mesenchymal tumors, and whether an EMT-equivalent process occurs, allowing for increased cellular adhesion, migration, and invasion, is unknown^53^. Still, given the similarities in the phenotypes of Tau-KD and EWSR1::FLI1-KD cells, we may hypothesize that Tau expression may participate in driving the morphologic and migration/invasion characteristics associated to EWSR1::FLI1 expression, which would suggest a key role of Tau in the metastatic processes highly predominant in EwS. It is therefore plausible that Tau may act on survival through the modulation of these pathways.

## Materials and methods

### Data and code availability

Cancer cell RNAseq data were downloaded from https://depmap.org/portal/. RNAseq data from PDXs were downloaded from https://www.cbioportal.org/. Gene expression microarray and proteomic data associated to the 18 EwS cell lines were already published^32^ and are deposited under accession code: GSE176190. Publicly available and pre-processed ChIP-Seq data from EwS cells corresponding to EWSR1::ETS and H3K27Ac were retrieved, analyzed and displayed using R2 genomics analysis and visualization platform (http://r2platform.com/escla/)^32^ for the *MAPT* locus. Single cell data of MSCs and EwS PDXs is publicly available under accession code GSE130025^34^. Any additional information required to reanalyze the data reported in this work paper is available upon request.

### Survival analysis

Kaplan-Meier survival analyses of mRNA expression data were carried out in 166 EwS patients whose molecularly confirmed and retrospectively collected primary tumors were profiled at the mRNA level by gene expression microarrays in previous studies^42^. To that end, microarray data generated on Affymetrix HG-U133Plus2.0, Affymetrix HuEx-1.0-st or Amersham/GE Healthcare CodeLink microarrays of the 166 EwS tumors (Gene Expression Omnibus (GEO) accession codes: GSE63157, GSE12102, GSE17618, GSE34620 provided with clinical annotations were normalized separately as previously described^54^. Only genes that were represented on all microarray platforms were kept for further analysis. Batch effects were removed using the ComBat algorithm^55^. Data processing was done in R. The validation cohort was previously described^42^. In Kaplan-Meier survival analyses, statistical differences between the groups were assessed by both the Mantel-Cox test.

### Immunohistochemistry

For the detection of Tau by IHC on xenografts and EwS patients’ tumors, 4-μm paraffin sections were incubated 1 h at 65°C and rehydrated using Ottix Plus and Ottix Shaper solutions according to manufacturer’s instructions (DiaPath X0076, X0096). Sections were subsequently washed 10 min in PBS 1X and incubated for 5 min at RT in 3% H_2_O_2_ to block peroxidase activity. Following two washes in PBS 1×, an antigen retrieval step was performed through incubation of the sections in 10 mM citrate buffer (pH 6) in a microwave (5 min at 730 W, 5 min at 400 W, 5 min at 520 W). After cooling down and two washes in PBS, blocking was achieved through a 1-h incubation at RT with normal horse serum (VECTASTAIN Elite ABC Universal PLUS Kit, Vector laboratories PK8200). The anti-Tau antibody Tau 13 (sc-21796, SantCruz Biotechnology) was then incubated at 2 μg/ml for 1 h at RT and the sections was then washed two times in PBS and incubated with biotinylated secondary antibodies for 1 h at RT (VECTASTAIN Elite ABC Universal PLUS Kit, Vector laboratories PK8200).Following two washes in PBS, the section was incubated in ABC solution (VECTASTAIN Elite ABC Universal PLUS Kit, Vector laboratories PK8200) for 30 min at RT, washed two times and incubated with DAB (DAB substrate kit, Vector laboratories SK-4100) until the section has become brown. The reaction was stopped washing the section in ddH_2_O, the tissues were counterstained with hematoxylin, washed and dehydrated using Ottix Shaper and Ottix Plus according to manufacturer’s instructions. The section was then mounted with glycerol. Images were acquired with a Zeiss Axio Lab.A1 Microscope (AxioCam ERc 5 s, Oberkochen, Germany). The intensity of marker immune reactivity was determined (grade 0=none, grade 1=low, grade 2=moderate and grade 3=strong).

### Generation of pseudolentiviral particles

To generate the inducible Tau-KD, we used an inducible lentiviral plasmid system: pLKO-Tet-On all-in-one vector (Addgene, 21915) containing a tetracycline-regulated promotor driving the expression of shRNA. The shRNA (shRNA-75) cloned in the inducible vector target the third repeat of the microtubule domain of Tau (5’- CCGGGTGTGGCTCATTAGGCAACATCTCGAGATGTTGCCTAATGAGCCACACTTTTTG-3’). For the constitutive downregulation of Tau, the Tau-specific shRNA 3127 (5’-GATCCAGCAGACGATGTCAACCTTGTGCTTCCTGTCAGACACAAGGTTGACATCGTCTGCCTTTTTG-3’) was inserted in the pGreenPuro vector (SI505A-1, System Biosciences). Pseudo-lentiviral particles were produced in HEK-293 cells (HEK 293TN, System Biosciences) with the pPACKH1 kit (LV500A-1, System Biosciences) by the calcium phosphate transfection method. Particles were harvested 48–72 h later, concentrated on a centrifugal filter (MWCO 30 kDa, UFC903024, Amicon), aliquoted and stored at −80 °C until use.

### Cell culture and lentiviral transduction

EwS cell lines TC-32 and TC-71 originated from the Childhood Cancer Repository (https://www.cccells.org). Cells are cultured in RPMI 1640 medium (ThermoFisher Scientific, 11875093) supplemented with 10% Fetal Bovine Serum without Tetracycline (FBS, 10270106, Gibco), 1% Penicillin-Streptomycin (PS, Gibco, 15140122), 1% Non-Essential Amino acids (NEAA, Gibco, 11140035). Cells were grown at 37 °C in saturated humidity and 5% CO2 and maintained in culture for less than one month.

For the generation of Tau-KD cells, parental TC-32 and TC-71 cells (1×10^5^) were seeded into a 24-well plate coated with poly-D-lysine (P6407, Sigma-Aldrich) one day before pseudo-lentiviral particle transduction. One day after transduction, cells were supplemented with fresh complete medium and selected in the presence of 2.5 μg/mL puromycin (P8833, Sigma-Aldrich) for two weeks. To induce the downregulation of Tau expression, TC-32 cells were incubated in the presence or absence of 0.5 μg/mL doxycycline (D9891, Sigma-Aldrich) for at least 2 weeks.

### Western blot

Cells (7×10^5^) were seeded into 6-well plates coated with poly-D-lysine (P6407, Sigma-Aldrich) in presence or absence of doxycycline. Two days after plating, total lysates were prepared in 100 μL of SDS-PAGE sample buffer (1.5% SDS, 8.3% glycerol, 0.005% bromophenol blue, 1.6% β-mercaptoethanol and 62.5 mM Tris pH 6.8) and incubated for 10 min at 100°C. 5 or 15 μL of the sample per lane was loaded on 10% SDS-polyacrylamide gels (SDS-PAGE). After SDS-PAGE, PVDF membranes with transferred proteins were incubated with the primary antibodies indicated in the figures: 0.2 μg/mL Tau13 (sc-21796 AF680, Santa-Cruz Biotechnology), 0.25 μg/mL rabbit monoclonal vimentin (ab92547, Abcam), 0.2 μg/mL FAK (ab76496, Abcam), 0.25 μg/ml P-FAK (611723, BD Transduction Lab) or 0.5 μg/mL GAPDH (ab181602, Abcam) antibody. Primary antibodies were revealed with anti-mouse IgG coupled to IRDye 680RD or anti-rabbit IgG coupled to IRDye 800CW (Licor Biosciences, 926–68070 & 926–32211) on a dual infrared imaging scanner (Licor Biosciences, Odyssey CLx 9140). Immune reactive bands were quantified with the software provided (Licor Biosciences, Image Studio V5.0.21, 9140–500).

### Immunostaining

For immune staining, cells (5×10^4^) were grown on poly-D-lysine coated 8-well microscope slides (80826-IBI, Ibidi). Cells were fixed in 4% paraformaldehyde and stained^13^ with primary antibodies: 2 μg/mL P-FAK antibody (ab81298, Abcam), 0.2 μg/mL paxillin antibody (AHO0492, Invitrogen). Detection by fluorescent laser confocal microscopy (Nikon C2 microscope) was done with 2 μg/mL secondary antibody anti-rabbit IgG-Alexa 647 (A21245, Thermo Fisher Scientific). Nuclei were counterstained with 0.5 μg/mL DAPI (D9542, Sigma-Aldrich). Phalloidin-iFluor488 (ab176753, Abcam) was incubated for 1 hour at RT according to manufacturer’s instruction. Images were acquired by sequential excitations (line-by-line scan) with the 405 nm laser (464/40 emission filter), the 488 nm laser (525/50 nm filter), the 561 nm laser (561/LP nm filter) and the 650 nm laser (594/633 emission filter). ImageJ was used for all image quantifications.

### Cell adhesion assay

96-well plates were coated with fibronectin (F4759, Sigma-Aldrich), laminin (L2020, Sigma-Aldrich) or Matrigel (CLS356234-1EA, Sigma-Aldrich) for 1 h at 37°C. Plates were washed twice with PBS 1X and RPMI complemented with 10% FBS for 1 h at 37°C, blocked with 1% BSA in 1X PBS for 1 h at RT. Following blocking, 5×10^4^ cells/well were added to plates in replicates of 4 to 6 per group. Plates were incubated at 37 °C for 30 min. Plates were washed with complete medium and fixed with methanol prior to staining with 0.05% crystal violet (HT90132, Sigma-Aldrich) in 20% ethanol for 20 min. Plates were washed with water and the dye was solubilized in 100% methanol. Absorbance per well was measured at 590 nm on a Tecan microplate reader.

### IncuCyte cell growth and wound healing assay

Cell growth was measured using IncuCyte Zoom (Essen Biosciences) Live-Cell Analysis Platform. 5×10^3^ TC-32 or 1×10^4^ TC-71 cells were plated in 6 replicates in a 96-well plate and were imaged every 2 h with a 4× objective. The percent confluence of each well at each time point was calculated using IncuCyte Zoom image processing software. All replicate values were obtained over an incubation time of 3 d and normalized over the cell density at DIV 0 for TC-32 and DIV1 for TC-71. To measure cell migration, TC-71 cells were plated at a density of 7×10^5^ cells/well. After 2 d, an open wound area was created in the cell monolayer using the IncuCyte® Wound Maker tool, washed with PBS and subsequently incubated with complete medium and wound closure was followed through image acquisition every 2 h.

### Transwell migration and invasion assays

Transwell migration assays were performed in Transwell 8 μm pore size (3422-48, Chemie Brunschwig AG)). TC-32 cells were seeded at 6×10^5^ cells/200 μL serum free-RPMI medium in the upper chamber. The lower chamber was filled with RPMI supplemented with 10% FBS. After 48-h incubation, the chambers were removed and fixed in 4% PFA for 10 min at RT, then washed with PBS 1×. The cells on the upper surface were removed using cotton swabs. Cells on the lower surface that migrated through the membrane were stained with 1% crystal violet in 2% Ethanol for 20 min at RT, washed thoroughly with water and air dried. Images (five per replicate) acquired at 10× magnification were quantified using ImageJ. This protocol is also followed for invasion assays except for the addition of 80 µl Matrigel (CLS356234-1EA, Sigma-Aldrich) diluted to 1 mg/ml in serum-free media) to the top of each insert 2 h at 37°C prior to adding cells.

### Statistics and reproducibility

Statistical analysis was performed with GraphPad Prism version 8.4 using the method specified in the legend of each figure. Exact *p*-values are specified in the figures. All quantifications were performed based on at least three independent biological replicates. Sample size, number of replicates and how they are defined is specified in the figure legends. When indicated, western blots and microscopic images are shown as representative data.

## Acknowledgments

We thank the whole laboratory for support and advice during this study. The laboratory of TGPG thanks its technicians Nadine Gmelin, Stefanie Kutschmann, Sabrina Knoth, and Felina Zahnow for excellent technical assistance. We thank Prof. Daniel Baumhoer for providing histological samples.

## Author contributions

Conceptualization: CM, SP, PP, FCA, TGPG

Methodology: FCA, MZ, JL, CM

Investigation: FCA, MZ, JL, CM, ALB, MC, MS, WH, UD

Supervision: SP, PP, TGPG

Writing: SP, PP, FCA, TGPG, MZ, JL

## Data and materials availability

all data generated or analyzed during this study are included in this published article and its supplementary information files.

## Supplementary Figure Legends

**Supplementary Figure 1.** Effect of doxycycline-treatment of parental TC-32 cells on adhesion, migration, and invasion. **A.** Lysates from biological triplicates were prepared from parental TC-32 cells not (ctrl) or treated (mock) with doxycycline for 2.5 weeks. Samples were analyzed by western blot with mouse Tau13 and rabbit GAPDH antibodies, followed by visualization with anti-mouse IgG IRDye 680RD and anti-rabbit IgG IRDye 800CW. Intensity of the signals is reported as ratio Tau/GAPDH (n=9). **B.** Parental TC-32 cells in MW96 were imaged for surface covered at the Incucyte every 2 h and cell proliferation calculated over 72 h starting from time 0 (n=12). **C.** Adherent parental TC-32 cells on the indicated cell substrates 30 min after plating and stained with crystal violet (OD590 nm normalized over ctrl (n= 4). **D.** TC-32 Tau-KD cells were seeded in transwell inserts without any coating (migration) or pre-coated with Matrigel (invasion). Cells were imaged on the other side of the filters 2 d later upon staining with crystal violet and quantified with ImageJ (n=3). Mock data were normalized over ctrl, reported as mean ± SD and analyzed with the Mann-Whitney test.

**Supplementary Figure 2.** Changes in P-FAK and vimentin are not artefacts of doxycycline treatment. Lysates from biological triplicates were prepared from parental TC-32 not (ctrl) or treated (mock) or with doxycycline for 2.5 weeks. Samples were resolved on SDS-PAGE and analyzed by western blot with mouse P-FAK and rabbit FAK antibodies, the rabbit vimentin antibody, and the rabbit GAPDH antibody. Primary antibodies were detected using anti-rabbit IgG IRDye 800CW or anti-mouse IgG IRDye 680RD. Signal intensity was quantified and reported as ratio P-FAK/FAK or vimentin/GAPDH (n=9). Mock data were normalized over ctrl, reported as mean ± SD and analyzed with the Mann-Whitney test.

