## Supplementary figures and images for "High Tau Expression Correlates with Reduced Invasion and Prolonged Survival in Ewing Sarcoma"

### Supplementary Figure 1

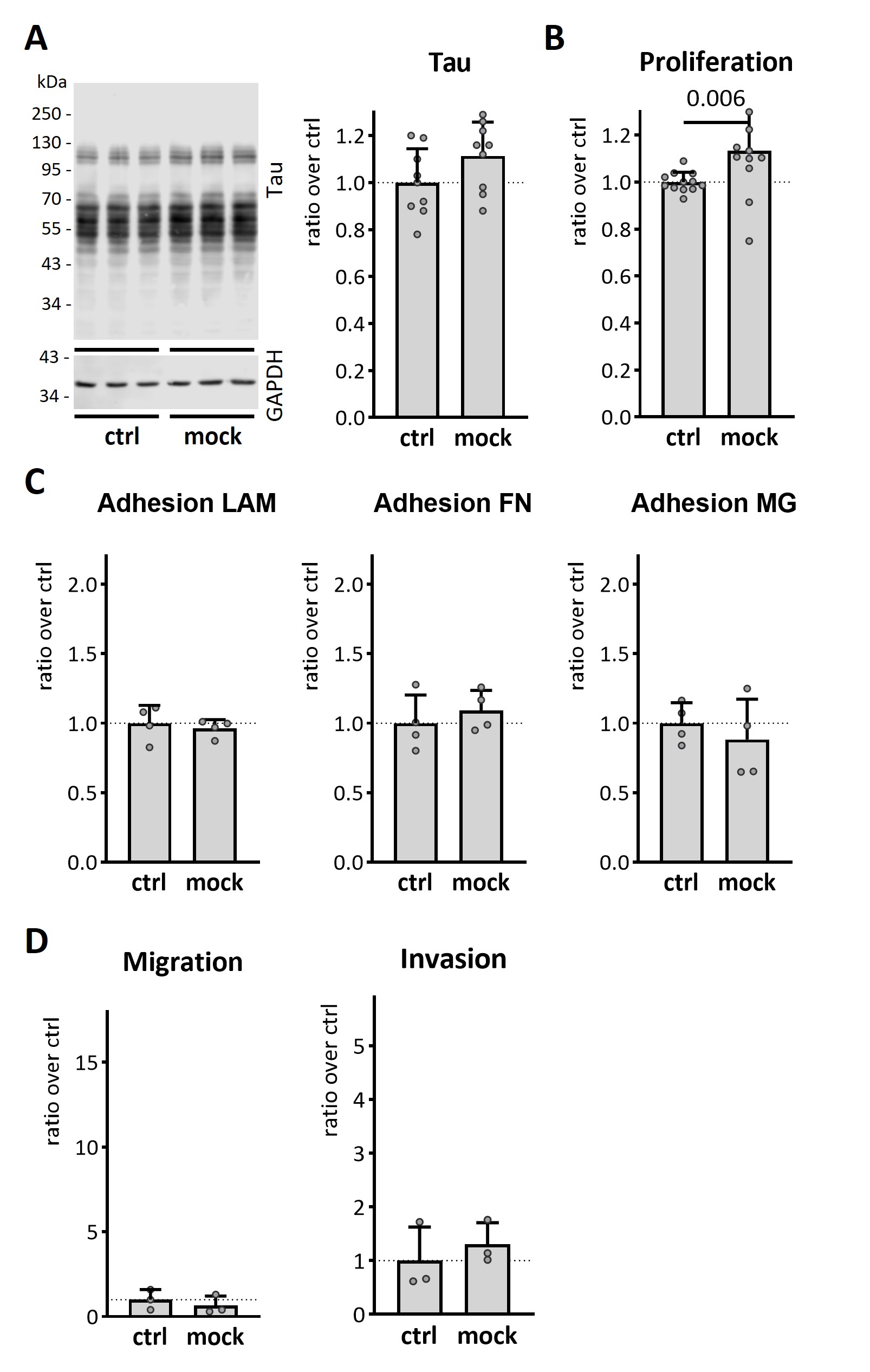

### Supplementary Figure 2

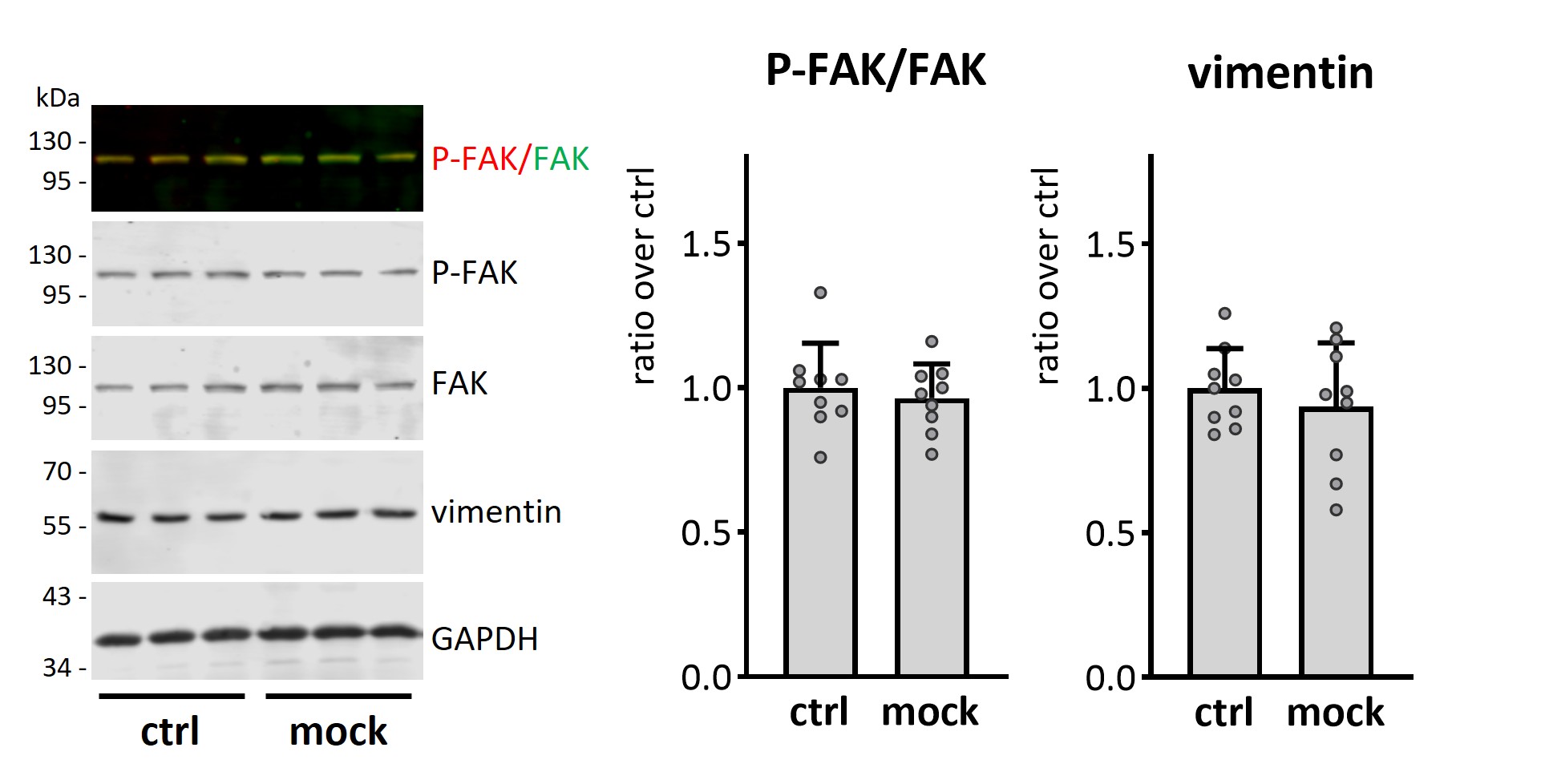
